# Mercury methylating microbial communities of boreal forest soils

**DOI:** 10.1101/299248

**Authors:** Jingying Xu, Moritz Buck, Karin Eklöf, Omneya Osman, Jeffra K. Schaefer, Kevin Bishop, Erik Björn, Ulf Skyllberg, Stefan Bertilsson, Andrea G. Bravo

## Abstract

The formation of the potent neurotoxic methylmercury (MeHg) is a microbially mediated process that has raised much concern because MeHg poses threats to wildlife and human health. Since boreal forest soils can be a source of MeHg in aquatic networks, it is crucial to understand the biogeochemical processes involved in the formation of this pollutant. High-throughput sequencing of 16S rRNA and the mercury methyltransferase, *hgcA*, combined with geochemical characterisation of soils, were used to determine the microbial populations contributing to MeHg formation in forest soils across Sweden. The *hgcA* sequences obtained were distributed among diverse clades, including *Proteobacteria, Firmicutes*, and *Methanomicrobia,* with *Deltaproteobacteria*, particularly *Geobacteraceae*, dominating the libraries across all soils examined. Our results also suggest that MeHg formation is linked to the composition of also non-mercury methylating bacterial communities, likely providing growth substrate (e.g. acetate) for the *hgcA*-carrying microorganisms responsible for the actual methylation process. While previous research focused on mercury methylating microbial communities of wetlands, this study provides some first insights into the diversity of mercury methylating microorganisms in boreal forest soils.

**Importance:** Despite a global state of awareness that mercury, and methylmercury in particular, is a neurotoxin that millions of people continue to be exposed to, there are sizable gaps in our fundamental understanding of the processes and organisms involved in methylmercury formation. In the present study we shed light on the diversity of the microorganisms responsible for methylmercury formation in boreal forest soils. All the microorganisms identified have a relevant role on the processing of organic matter in soils. Moreover, our results show that the formation of methylation formation is not only linked to mercury methylating microorganisms but also to the presence of non-mercury methylating bacterial communities that contribute to methylmercury formation by the appropriate substrate to the microorganisms responsible for the actual methylation process. This study improves current knowledge on the diversity of organisms involved in methylmercury formation in soils.

## INTRODUCTION

Mercury (Hg) is a potent toxin that might cause severe negative effects on wildlife and human health (1). The toxicity of Hg is of such concern that 128 countries have signed the Minamata Convention, a global treaty that entered into force in August 2017 with the explicit objective to reduce Hg emissions and protect human health and the environment. High Hg emissions in the past have led to high present-day Hg levels in different parts of the atmosphere, oceans and terrestrial ecosystems (2, 3). Because Hg has a strong affinity for reduced sulphur or thiol (RSH) functional groups of soil organic matter (OM) (4, 5), the increased atmospheric deposition of Hg during the industrialisation has resulted in high Hg concentrations in organic-rich soils (6). As a consequence, the typically OM-rich soils in the boreal biome has retained Hg deposition from both natural and anthropogenic emissions, and now represent an important global Hg stock (4, 7).

Soil OM has also been identified as a main vector of Hg and methylmercury (MeHg) transport from catchments to surface waters in boreal areas (8, 9). Indeed, the mobilisation of inorganic Hg (Hg^(II)^) and, the more harmful, MeHg from soils by means of OM-mediated transport has been linked to MeHg accumulation in lake sediments within catchments (9, 10) and in fish (11). As high MeHg levels in fish have raised much concern in many boreal regions over the past decades (12, 13) and since forest soils are an important site for MeHg formation (14), it is crucial to understand the processes and the organisms involved in MeHg formation in boreal soils.

The methylation of Hg^(II)^ to MeHg is biologically mediated (15) and takes place under oxygen deficient conditions typical for wetlands (16), water logged soils (14), sediments (9) and anoxic water columns (17), but can also occur in suspended particles in the aerobic zone of aquatic systems (18, 19). Specific strains of sulphate-reducing bacteria (20, 21), iron reducing bacteria (FeRB) (22, 23), methanogens (24) and Firmicutes (25) have the capability to methylate Hg^(II)^. However, a number of factors controlling bacterial activity and/or the geochemical speciation of inorganic Hg^(II)^ will govern MeHg formation in the environment (9, 26). For example, increases in temperature might lead to increases in biological activity and accordingly also higher Hg^(II)^ methylation rates (27). Redox potential also seems to be a key factor as suboxic and mildly reducing conditions seem to promote high Hg^(II)^ methylation rates, whereas anoxic and strongly reducing conditions might lead to elevated sulphide concentrations that eventually prevent Hg^(II)^ from being available for methylation (28). Sulphur plays a major role in influencing Hg^(II)^ methylation by directly affecting the activity of some methylating bacteria (e.g. sulphate reducing bacteria, SRB) and/or control the availability of Hg^(II)^ for methylation (5). Specific organic matter (OM) compounds can promote Hg^(II)^ methylation by enhancing bacterial activity (9), but also by defining Hg^(II)^ speciation (29) and Hg^(II)^ availability (30, 31). OM can also facilitate Hg^(II)^ methylation by inhibiting mercury sulphide (HgS(s)) precipitation or enhance HgS(s) dissolution thereby providing available Hg^(II)^ for methylating microorganisms (32). High OM concentrations might also decrease Hg methylation by formation of high mass molecular mass complexes that hamper Hg^(II)^availability (30). Recently it has been concluded that the availability of Hg^(II)^ depends heavily on the S^(-II)^ concentration in porewater and the RSH(aq)/RSH(ads) molar ratio of DOM (29). Together, all these studies highlight that geochemical conditions are key in determining the availability of Hg^(II)^ and the activity of the microbial communities involved in the process.

The identification of two functional genes, *hgcA* and *hgcB*, which play essential roles in Hg^(II)^ methylation (15), provided the means to more directly characterise the complexity of microbial communities involved in the formation of MeHg in natural ecosystems. This approach has been applied to marshes, sediments and swamps in several geographic regions (33–36); rice paddies in China (37), and water conservation areas of the northern Everglades, USA (38). However, very little work to date has been conducted to reveal the distribution of microbial groups responsible for Hg^(II)^ methylation in forest soils within the vast boreal biome. To the best of our knowledge, no studies have directly described the composition and spatial variation in Hg^(II)^ methylating microbial communities in such forests. Therefore, the primary goal of this paper was to describe Hg^(II)^ methylating microbial communities in various boreal forest soils and identify soil characteristics important for shaping these communities. High-throughput next generation sequencing of amplified 16S rRNA and *hgcA* genes combined with molecular barcoding and detailed soil geochemical characterisations were performed to study the Hg^(II)^ methylating microbial communities in 200 soil samples from three different boreal forest regions in order to shed light on the biogeography of microorganisms responsible for MeHg formation in the boreal landscape.

## RESULTS

### Bacterial community composition in boreal forest soils

Soil samples were collected from 200 sites in October 2012 and were distributed across eight catchments in three boreal forest regions in Sweden (Table S1, Table S2). A total of 3 321 197 high quality 16S rRNA sequences remained after quality control and chimera removal (7–72 911 reads per sample). The sample with only 7 reads was removed, and we then rarefied the rest of the data to the remaining sample with the fewest reads (1692 reads). The final rarefied sequence dataset (329 940 reads) clustered into 33 158 operational taxonomic units (OTUs) using a similarity threshold of 97 %. In the rarefied dataset, 35 taxa at phyla level, 69 taxa at class level, 119 taxa at order level, and 187 taxa at family level were detected from all the soil samples across three regions. The overall coverage of the forest bacterial community is reflected in the combined richness detected for random subsets of analysed samples. The logarithmic shape indicated that most of the considerable OTU richness occurring in the forest soils was accounted for in the combined dataset (Fig. S1). Among the dominant phyla across all regions (>5 % relative abundance), *Acidobacteria* was the most abundant, followed by *Proteobacteria, Planctomycetes, Bacteroidetes, Parcubacteria and Verrucomicrobia* (Table 1). Combined, these phyla accounted for 77.5 % of the total sequences (Table 1). Most of the previously identified clades known to contain Hg^(II)^ methylators (25, 39) were detected in the present study, including *Deltaproteobacteria* (3.31 % of the total reads)*, Chloroflexi* (2.60 % of the total reads), *Firmicutes* (0.77 % of the total reads) and *Euryarchaeota* (0.66 % of the total reads) (Table 1). Microbial community composition based on 16S rRNA sequences in the 34 studied MeHg hotspots showed a similar pattern in terms of the dominant phyla (>5 % relative abundance), with *Acidobacteria* and *Proteobacteria* being the most abundant ones. However, *Bacteroidetes* and *Chloroflexi* contributed much more to the total communities at these hotspots compared to the combined dataset across all 200 samples (Table 1).

**Table 1.** Comparison of the relative abundances (%) of the most abundant taxa (phylum level) in all the samples (n=200) with the 34 MeHg hotspots based on 16S rRNA sequences. Relative abundances of classes under phylum *Proteobacteria* are listed with indent (SD: Standard deviation)

| Most abundant taxa | Mean ± SD |  | Maximum |  | Minimum |  |
| --- | --- | --- | --- | --- | --- | --- |
|  | All samples | Hotspots | All samples | Hotspots | All samples | Hotspots |
| *Acidobacteria* | 36.11 ±10.53 | 25.57 ±8.77 | 73.64 | 49.29 | 8.10 | 9.40 |
| *Proteobacteria* | 13.99 ±4.03 | 16.56 ±2.96 | 28.13 | 27.60 | 2.90 | 8.87 |
| *Alphaproteobacteria* | 6.83 ±3.01 | 7.13 ±2.81 | 16.43 | 13.95 | 1.77 | 2.66 |
| *Deltaproteobacteria* | 3.31 ±1.69 | 3.56 ±1.38 | 13.36 | 7.15 | 0.71 | 1.30 |
| *Gammaproteobacteria* | 2.06 ±1.33 | 1.48 ±0.76 | 7.15 | 3.66 | 0.24 | 0.35 |
| *Betaproteobacteria* | 1.78 ±2.13 | 4.14 ±2.47 | 11.11 | 10.46 | 0.00 | 0.65 |
| *Epsilonproteobacteria* | 0.01 ±0.03 | 0.03 ±0.06 | 0.30 | 0.24 | 0.00 | 0.00 |
| *Planctomycetes* | 8.18 ±4.21 | 5.82 ±2.77 | 24.82 | 11.64 | 1.36 | 1.95 |
| *Bacteroidetes* | 6.61 ±5.24 | 11.38 ±7.92 | 51.60 | 51.60 | 0.41 | 1.60 |
| *Parcubacteria* | 6.35 ±4.19 | 9.01 ±5.14 | 26.36 | 24.47 | 0.06 | 2.13 |
| *Verrucomicrobia* | 6.28 ±2.78 | 5.30 ±2.31 | 14.89 | 10.64 | 0.65 | 0.65 |
| *Thaumarchaeota* | 3.96 ±2.77 | 2.53 ±2.44 | 18.44 | 14.83 | 0.00 | 0.00 |
| *Actinobacteria* | 3.11 ±2.38 | 2.94 ±1.62 | 19.86 | 6.03 | 0.47 | 0.89 |
| *Chlamydiae* | 2.83 ±2.56 | 1.31 ±1.08 | 22.87 | 3.71 | 0.24 | 0.30 |
| *Chloroflexi* | 2.60 ±3.18 | 7.16 ±5.18 | 17.79 | 15.19 | 0.00 | 0.12 |
| *Others* | 9.97 ±0.89 | 12.41 ±1.66 | 17.14 | 8.98 | 0.00 | 0.00 |

A non-metric multidimensional scaling (nMDS) plot based on 16S sequences was used to visualise the composition of the bacterial community among samples. *Unclassified Acidobacteriales, Unclassified Ignavibacteriales, Spirochaetaceae, Holophagaceae*, *Anaerolineaceae, Betaproteobacteria and Tepisiphaeraceae* were important contributing families for shaping the differences in bacterial community composition among samples (Fig. 1). Geochemical factors that were correlated (correlation coefficients > 0.5) with the bacterial composition were projected on top with longer vectors implying stronger correlations (Fig. 1). %MeHg, reflected by bubble sizes, presented a strong coupling to the bacterial community composition, which was further confirmed by %MeHg presenting a long vector among all the geochemical factors (Fig. 1). Water content, C%, S% and N% were all found to be the factors that affected the composition of soil bacterial community (Fig. 1), indicating that a supply of organic matter and nutrients in the moist soil shapes the bacterial community. This is in agreement with previous research that pointed out the contribution of nutrients and organic matter to bacterial activities and Hg^(II)^ methylation (9, 28). Also, S was well correlated with both C and N (Table S3), suggesting that most of the measured sulphur in the sampled soils has likely an organic origin. This has been found as a common feature in boreal soils (40–42).

**Figure 1.**
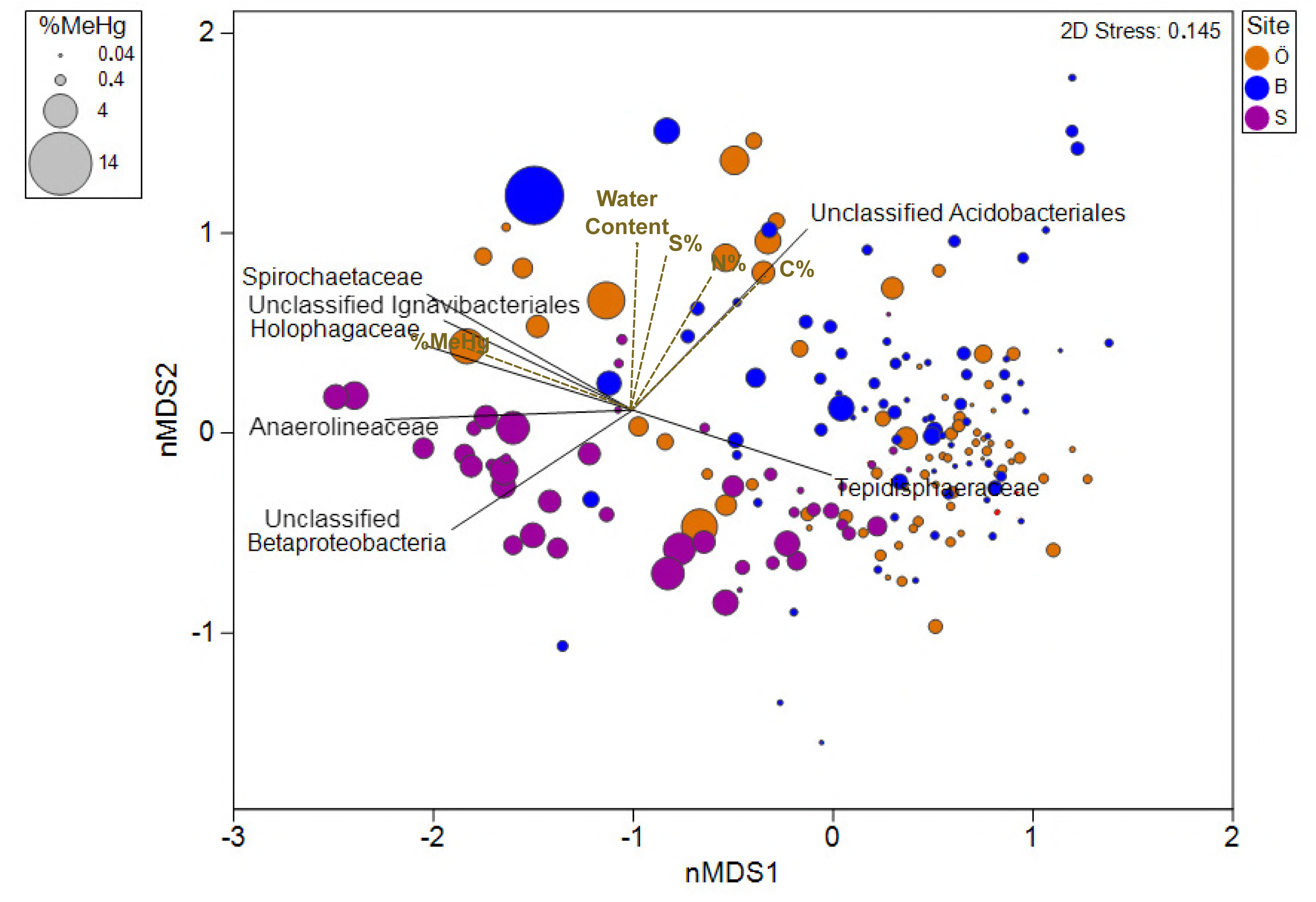
Non-metric multidimensional scaling (nMDS) of microbial community composition of all samples (family level based on 16S rRNA) overlaid with families (black lint) and geochemical factors (dotted brown line) moderately correlated with biotic ordination (correlation coefficients > 0.5) (%MeHg: MeHg/THg). Relative dissimilarities (or distances) among the samples were computed according to the resemblance matrix calculated on fourth rooted family reads.

*Unclassified Fibrobacterales, Methanosaetaceae, Unclassified Ignavibacteriales, Spirochaetaceae, Holophagaceae* and *Anaerolineaceae* exhibited the highest correlations with %MeHg (Table 2). *Syntrophobacteraceae, Methanosarcinaceae, Methanoregulaceae, Desulfobulbaceae, Syntrophaceae, Desulfobacteraceae* and *Dehalococcoidaceae*, which potentially host Hg^(II)^ methylators (25, 39), were also found relevant to the bacterial community composition in high-%MeHg sites (Table 2).

**Table 2.**
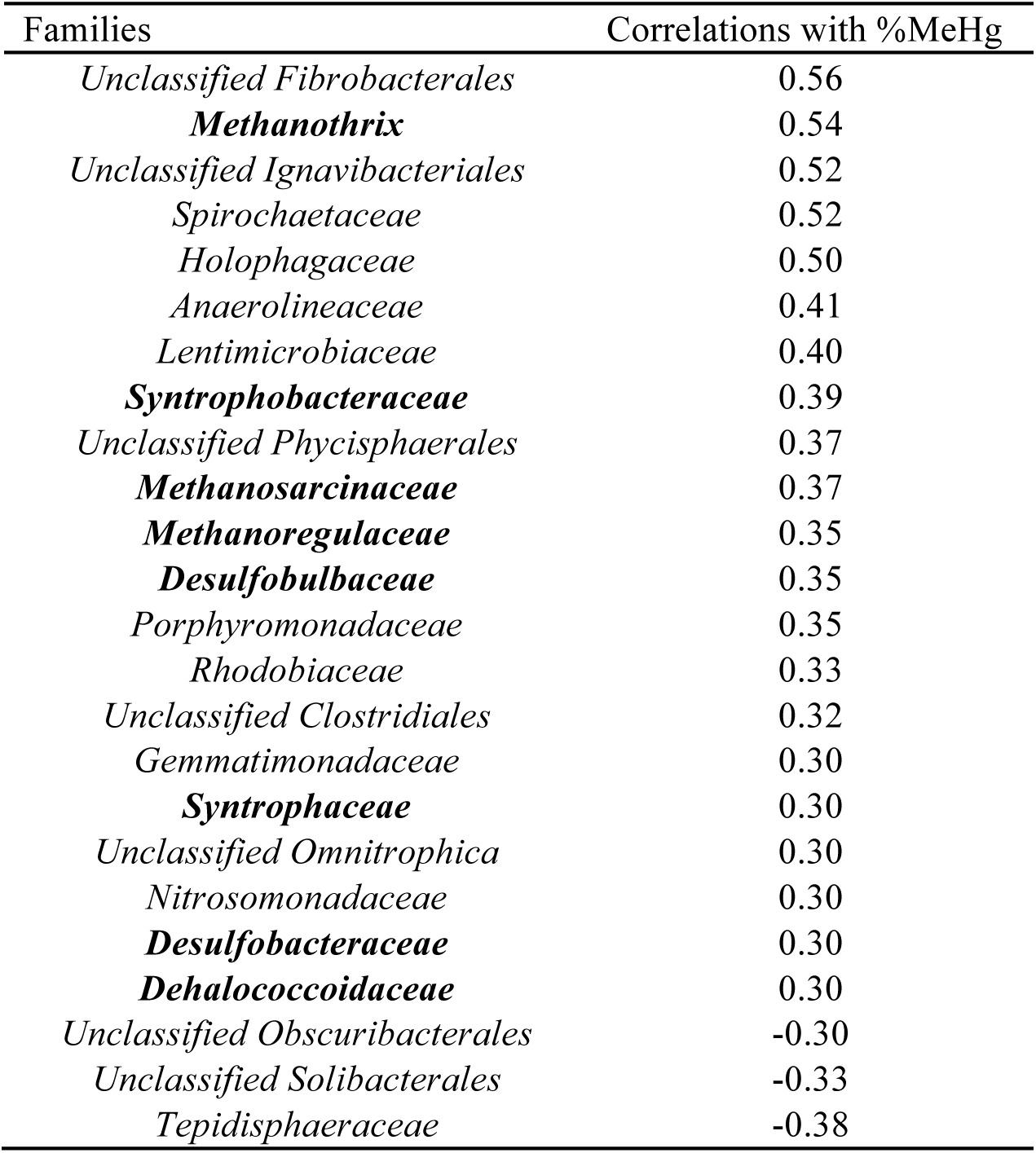
Moderate (0.5≤R<0.7) to weak (0.3≤R<0.5) Pearson correlations between families and %MeHg in all samples based on 16S rRNA. Families potentially involved in Hg methylation were marked in bold.

| Families | Correlations with %MeHg |
| --- | --- |
| *Unclassified Fibrobacterales* | 0.56 |
| ***Methanotherix*** | 0.54 |
| *Unclassified Ignavibacteriales* | 0.52 |
| *Spirochaetaceae* | 0.52 |
| *Holophagaceae* | 0.50 |
| *Anaerolineaceae* | 0.41 |
| *Lentimicrobiaceae* | 0.40 |
| ***Syntrophobacteraceae*** | 0.39 |
| *Unclassified Phycisphaerales* | 0.37 |
| ***Methanosarcinaceae*** | 0.37 |
| ***Methanoregulaceae*** | 0.35 |
| ***Desulfobulbaceae*** | 0.35 |
| *Porphyromonadaceae* | 0.35 |
| *Rhodobiaceae* | 0.33 |
| *Unclassified Clostridiales* | 0.32 |
| *Gemmatimonadaceae* | 0.30 |
| ***Syntrophaceae*** | 0.30 |
| *Unclassified Omnitrophica* | 0.30 |
| *Nitrosomonadaceae* | 0.30 |
| ***Desulfobacteraceae*** | 0.30 |
| ***Dehalococcoidaceae*** | 0.30 |
| *Unclassified Obscuribacterales* | -0.30 |
| *Unclassified Solibacterales* | -0.33 |
| *Tepidisphaeraceae* | -0.38 |

### Distribution of Hg^(II)^ methylators

The samples with high soil MeHg concentrations and %MeHg > 1% were defined as “MeHg hotspots”. In 34 MeHg hotspots (see soils geochemistry descriptors in Table S4, n = 34), the relative abundance of microbial families carrying representatives known to methylate Hg^(II)^ was assessed based on *hgcA* sequences (25, 39). A total of 1 257 577 *hgcA* sequences remained after quality control and chimera removal (11 404–55 461 reads per sample). The *hgcA* dataset was rarefied to the remaining sample with the fewest reads (11 404 reads). The rarefied sequence dataset accounted a total of 387 736 reads that clustered into 573 operational taxonomic units (OTUs) using a similarity threshold of 97 %. As for the 16 rRNA, the logarithmic shape indicated that most of the considerable species richness of Hg^(II)^ methylators occurring in the forest soils was accounted for in the combined dataset (Fig. S1). Representative sequences from 22 families were found in the 34 analysed MeHg hotspots. Of all the *hgcA* sequences, 3.13 % were not taxonomically assigned (Unclassified), 0.28 % were unclassified *Euryarchaeota*, and 7.28 % could not be assigned beyond the rank of Bacteria (Unclassified Bacteria). The majority of the sequences annotated to the level of family clustered with *Deltaproteobacteria*, making up 85.4 % of all the *hgcA* reads (Table 3). The remaining classified *hgcA* sequences were distributed across diverse families affiliated to *Firmicutes* and *Methanomicrobia*. *Unclassified Deltaproteobacteria* represented up to 56 % of the reads and among the identified families, *Geobacteraceae* were the most abundant, contributing up to 40 % in Strömsjöliden. *Ruminococcaceae* (3.21 % of all *hgcA* reads) occurred as another important family in the hotspots in Örebro; while methanogens and syntrophic lineages were less abundant in the hotspots based on *hgcA* sequences (Table 3).

**Table 3.** Relative abundance of families involved in Hg^(II)^ methylation based on *hgcA* sequences in 34 hotspots.

| Families | Örebro | Balsjö | Strömsjöleden |
| --- | --- | --- | --- |
|  | % of *hgcA* reads | % of *hgcA* reads | % of *hgcA* reads |
| *Unclassified Deltaproteobacteria* | 43.24±37.11 | 44.85±30.09 | 55.69±18.23 |
| *Geobacteraceae* | 26.79±31.09 | 24.62±22.22 | 39.40±18.96 |
| *Unclassified Bacteria* | 10.72±17.45 | 25.58±33.67 | 1.43±1.02 |
| *Ruminococcaceae* | 9.12±18.23 | 1.52±2.30 | 0.15±0.04 |
| *Unclassified* | 6.62±8.65 | 2.37±3.86 | 1.27±2.98 |
| *Unclassified Euryarchaeota* | 0.84±2.22 | 0.02±0.02 | 0.01±0.02 |
| *Desulfovibrionaceae* | 0.83±1.28 | 0.16±0.03 | 0.02±0.04 |
| *Unclassified Methanomicrobiales* | 0.49±1.21 | 0.06±0.09 | 0.03±0.12 |
| *Syntrophaceae* | 0.35±0.45 | 0.05±0.00 | 0.00±0.00 |
| *Methanomassiliicoccaceae* | 0.31±0.53 | 0.02±0.00 | 0.13±0.05 |
| *Methanoregulaceae* | 0.20±0.03 | 0.06±0.03 | 0.00±0.01 |
| *Syntrophomonadaceae* | 0.17±0.13 | 0.02±0.03 | 0.13±0.04 |
| *Unclassified Desulfovibrionales* | 0.14±0.15 | 0.02±0.05 | 0.03±0.04 |
| *Unclassified Clostridiales* | 0.06±0.22 | 0.51±0.19 | 0.08±0.07 |
| *Unclassified Firmicutes* | 0.06±0.02 | 0.00±0.00 | 0.00±0.00 |
| *Unclassified Desulfuromonadales* | 0.03±0.00 | 0.10±0.03 | 0.39±0.32 |
| *Desulfobulbaceae* | 0.02±0.02 | 0.01±0.00 | 0.00±0.01 |
| *Desulfuromonadaceae* | 0.01±0.01 | 0.00±0.04 | 0.00±0.04 |
| *Syntrophorhabdaceae* | 0.01±0.00 | 0.00±0.02 | 0.00±0.06 |
| *Unclassified Deferrisoma* | 0.01±0.00 | 0.00±0.00 | 1.18±0.98 |
| *Desulfarculaceae* | 0.00±0.02 | 0.00±0.00 | 0.00±0.02 |
| *Pelobacteraceae* | 0.00±0.01 | 0.01±0.01 | 0.07±0.03 |

*Unclassified Desulfuromonadales, Geobacteraceae, Ruminococcaceae, unclassified Desulfovibrionales, Desulfovibrionaceae, and unclassified Deltaproteobacteria* seemed to contribute to differences in the composition of Hg^(II)^ methylators in the studied soils (Fig. 2a). Among the measured geochemical parameters, the S% and the C/S seemed to have an impact on shaping the community composition of Hg^(II)^ methylators (Fig. 2b). Moreover, *Methanoregulaceae, Desulfovibrionaceae, Desulfuromonadaceae, Desulfarculaceae* and *Methanomassiliicoccaceae* correlated positively with S% and negatively with C/S (Table S5). In the studied MeHg hotspots, S was strongly correlated with both C and N (Table S6), suggesting most of the measured sulphur in the hotspots is also likely presented in organic forms.

**Figure 2.**
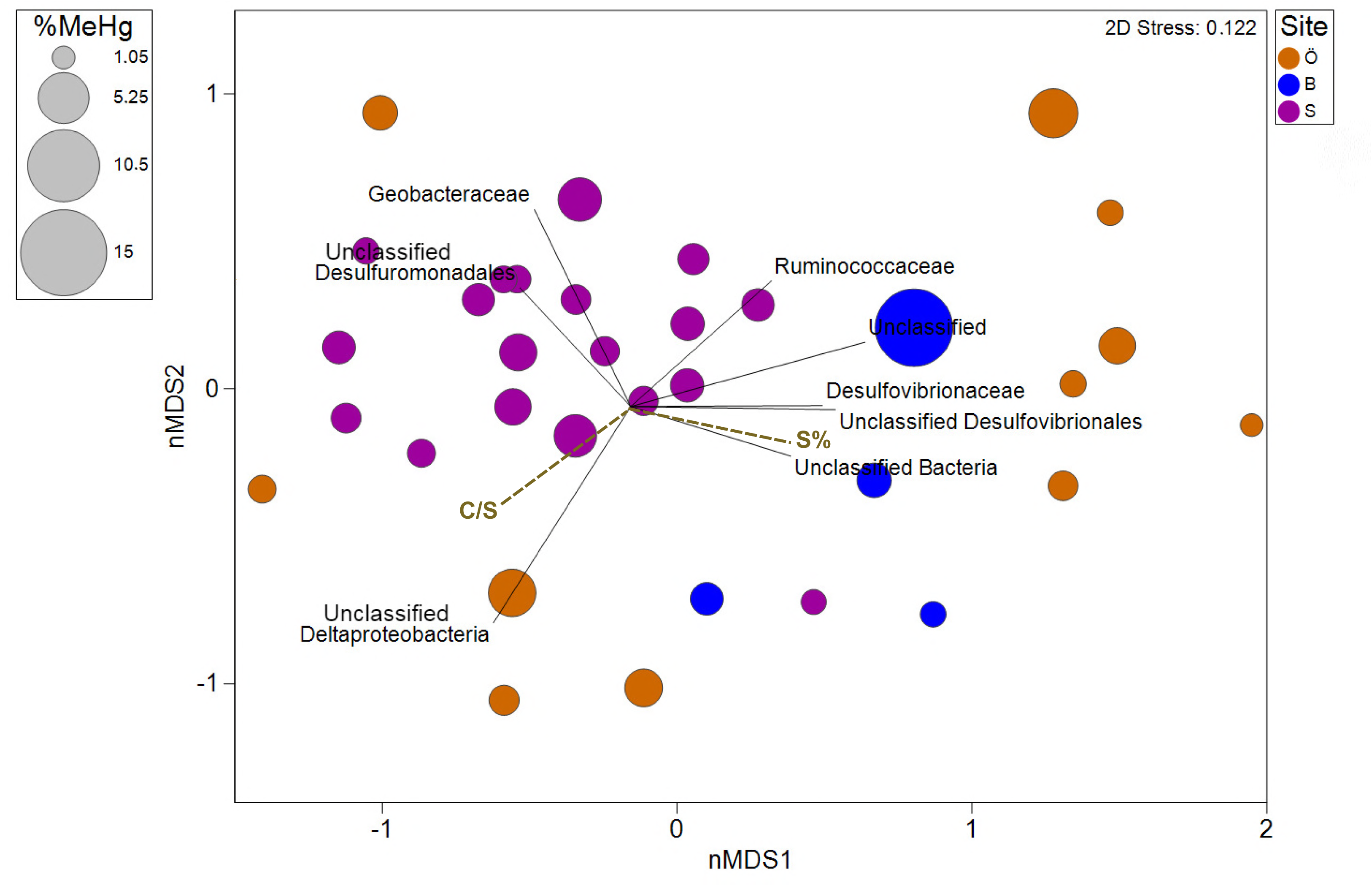
Non-metric multidimensional scaling (nMDS) of potential Hg methylators (family level based on *hgcA*) in 34 hotspots overlaid with geochemical factors that were moderately correlated with the biotic ordination positions (correlation coefficients > 0.5)

### Phylogenetic analysis of *hgcA* genes

All the *Proteobacteria* families belonged to *Deltaproteobacteria*, a class with which most currently confirmed Hg^(II)^-methylating bacteria are affiliated (43, 44). When combined, the 20 most abundant OTUs accounted for 72 % of the total reads. Noteworthy, phylogenetic analysis revealed that the most abundant Hg^(II)^-methylating OTUs (“OTU_0005”, “OTU_0705”, “OTU_0008”, and “OTU_0012”) in the studied forest soils were either taxonomically assigned as *Geobacter sp. or* phylogenetically related to *Geobacter* species (Fig. 3). Among the 20 most abundant OTUs, 17 were taxonomically annotated as *Deltaproteobacteria.* Among these 17 OTUs, 9 were taxonomically annotated as *Geobacter* and 8 were phylogenetically related to *Geobacter* species (Fig. 3). Summing the identified *Geobacter* and the OTUs phylogenetically related to *Geobacter* species, these 17 OTUs accounted for 62 % of the total *hgcA* reads. The 5^th^ most abundant OTU and was taxonomically denoted as *Firmicutes* (*Ethanoligenens*) and the 6^th^ and 7^th^ could not be annotated beyond the bacterial domain.

**Figure 3.**
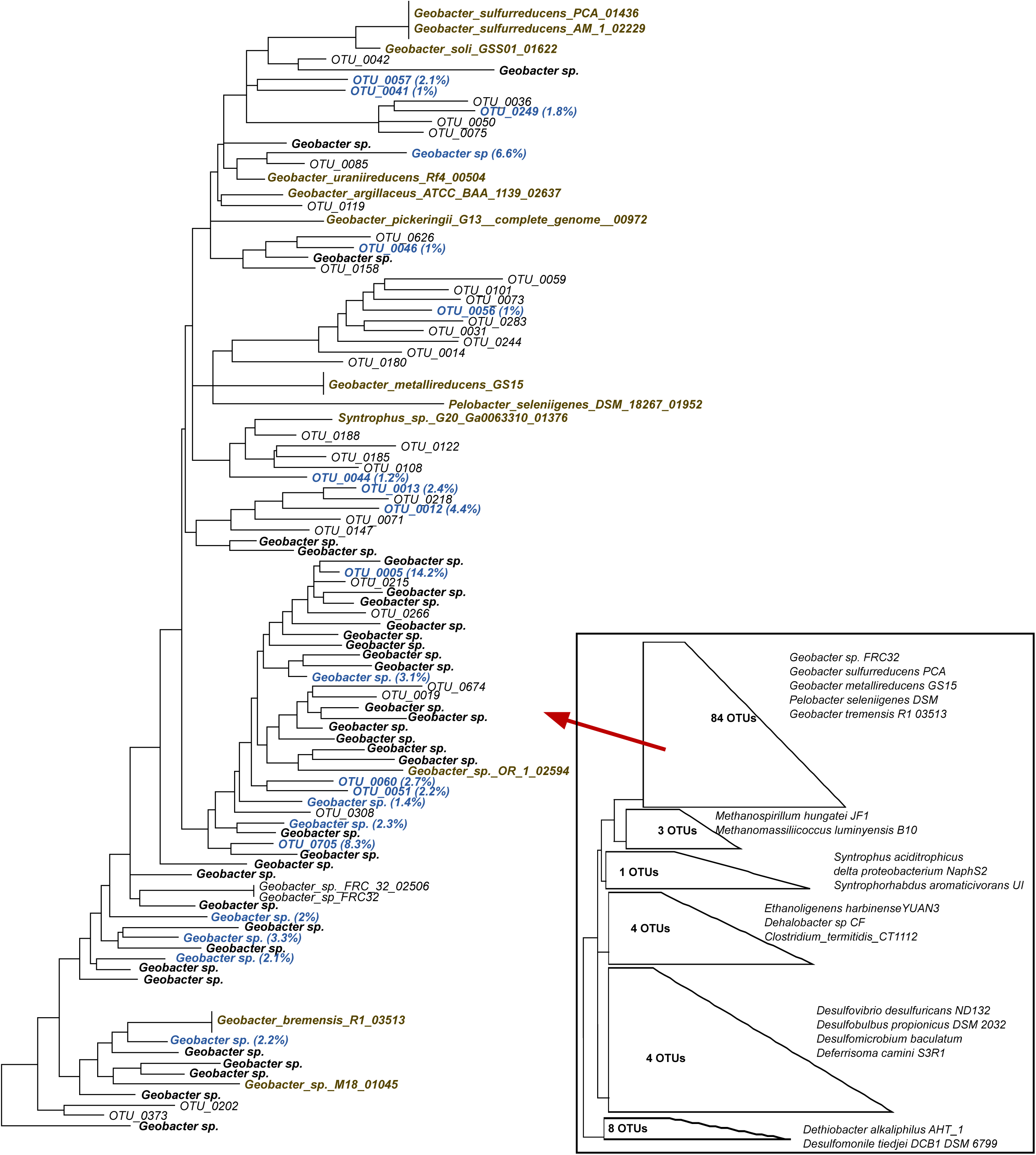
Phylogenetic relationships of *Deltaproteobacterial hgcA* sequences in the studied forest soils. The 20 most abundant *Deltaproteobacteria* are in blue. The OTUs taxonomically assigned as *Geobacter* are indicated in the plot “*Geobacter sp.”*. OTUs non-taxonomically assigned are presented as “OTU”. Reference genomes are marked in brown. The tree was generated using RAxML (version 8.2.4) with the PROTGAMMLG model and the autoMR to choose the number of necessary bootstraps (750). Please see details of the collapsed tree in the Fig. S2.

## DISCUSSION

### Community composition of Hg^(II)^ methylators in boreal forest soils

Among the diverse microbial communities seen in the soil samples (Table 1), most of the previously identified Hg^(II)^ methylating groups, e.g., *Deltaproteobacteria, Chloroflexi, Firmicutes and Euryarchaeota* could be detected (Table 3). *Deltaproteobacteria* have been considered a predominant Hg^(II)^ methylating class in anaerobic soils (34, 37, 38). In the present study, *Deltaproteobacteria* were also the predominant Hg^(II)^ methylators at the hotspots with *Geobacteraceae* as the most represented family. This family alone contributed over 30% of all *hgcA* reads, and their importance could be seen at all the sampled sites and particularly in Strömsjöliden (Table 3). Iron reducing bacteria (FeRB) have previously been shown to be important for Hg^(II)^ methylation in some environments (22, 23, 36, 43), and most *Geobacter* tested so far are particularly efficient at MeHg formation in the laboratory (23). This suggests that the ability to methylate Hg^(II)^ is widely distributed and a typical feature among the *Geobacteraceae*. The lack of a specific inhibitor for FeRB have hindered the quantification of the relative contribution of FeRB compared to SRB (i.e molybdate inhibitor) and methanogens (i.e. Bromoethanesulfonate inhibitor) to MeHg formation. The discovery of the *hgcA* pushed the state of the art and made possible to identify Hg^(II)^ methylators in environment (15, 25). Our results combined with previous findings in wetlands and paddy soils (34, 37, 38) highlight the importance of *Geobacteraceae* as Hg^(II)^ methylators in boreal forest soils and evidence their potentially very important roles in a wide range of environments.

While SRB are considered to be the principal Hg^(II)^ methylators in aquatic systems (27, 45–48), not much information is available on Hg^(II)^ methylators in soils. However, identified SRB in the hotspots only accounted for a minor portion of Hg^(II)^ methylators (Table 3). However, it is nevertheless plausible that at least some of the *hgcA* sequences annotated as unclassified *Deltaproteobacteria* (Table 3) could be unknown Hg^(II)^ methylating SRB or even Hg^(II)^ methylating sulphate-reducing syntrophs, capable of syntrophic fermentation of simple organic acids in the absence of sulphate as the terminal electron acceptor (49, 50). Therefore, we cannot discard the possibility that also SRB contribute significantly to Hg^(II)^ methylation in the studied systems. A previous study based on selective inhibitors and rate measurements indeed suggested SRB played an important role in MeHg formation in boreal forest soils (41). Additionally it has been demonstrated that even when SRB belong to the ‘rare biosphere’ of peatlands, they contribute significantly to respiration processes (51).

*Ruminococcaceae* belongs to another newly confirmed representative of Hg^(II)^ methylators, the *Firmicutes* (25). *Firmicutes* contributed to Hg^(II)^ methylating microbial communities at the water conservation areas of the Florida Everglades (38) but were not detected in boreal wetlands (34). In the present study, *Ruminococcaceae* were prominent contributors to the *hgcA* pool in hotspots from Örebro and in all soils from Strömsjöliden (Table 3). They could thus play a role in shaping the composition of Hg^(II)^ methylating community as further indicated by the negative correlation though weak between *Ruminococcaceae* and C/S, a primary geochemical factor shaping Hg^(II)^ methylating communities in the hotspots (Table S5 and Fig. 2b). Not much research has been devoted to the possible relationship between organic S and Hg^(II)^ methylating *Ruminococcaceae*. Considering the abundance of this group in forest soils, further efforts are needed to shed light on the metabolic or physiological pathways of Hg^(II)^ methylating *Ruminococcaceae*.

Methanogens were early on suspected to be responsible for Hg^(II)^ methylation (52), but not until recently were they verified as a significant source of Hg^(II)^ methylators in various environments (24, 34). In the hotspots in the studied soils, they were also detected, though not very abundant in the Hg^(II)^ methylating microbial community. *Chloroflexi* has recently been identified as potential Hg^(II)^ methylators in the water conservation areas, paddy soils and wetlands (34, 38, 53). The *hgcA* data did not confirm any significant role of this group in MeHg production in boreal forest soils (Table 3), even though 16S rRNA data revealed non-Hg^(II)^ methylating *Chloroflexi* (e.g. the class *Anaerolineae*) in soils from all three regions (Table 1).

Previous studies have mainly explored flooded environments such as paddy soils (37), boreal wetlands (34) and the water areas of the Florida Everglades (38). Hence our study provided important new information on the composition and diversity of Hg^(II)^ methylating microbial communities in non flooded boreal forest soils and the boreal landscape, and in doing so identified *Geobacteraceae* as significant Hg^(II)^ methylators in the terrestrial biome. The diversity of Hg^(II)^ methylators described in this study need to be interpreted cautiously. The *hgcA* gene was only recently discovered and the optimization of the appropriate methods and, in particular the design of primers for the *hgcA* amplification, is still developing (54). Additionally, DNA based methods only reveal the presence of organisms, while alternative approaches based on transcription data, proteomes or rate measurements are needed for verifying their activity. Our data nevertheless provide new insights about Hg^(II)^ methylating microbial communities in boreal forest soils and can as such guide and serve as a resource for future research efforts in this field.

### Bacterial communities fuel Hg^(II)^ methylators

%MeHg has previously been used as a proxy for methylation efficiency (55, 56), and high %MeHg has also in a few cases been shown to correlate positively with the abundance of Hg^(II)^ methylators (14, 57). In the current study, sites with high %MeHg featured bacterial communities different from those observed at sites with low % MeHg (Fig. 1). Although, families known to contain Hg^(II)^ methylators (*Syntrophobacteraceae, Methanosarcinaceae, Methanoregulaceae, Desulfobulbaceae, Syntrophaceae, Desulfobacteraceae* and *Dehalococcoidaceae*; 25) were found at sites with high %MeHg, there were also positive correlations between %MeHg and families that are not known to host Hg^(II)^ methylators, such us *Unclassified Fibrobacterales, Methanothrix* (formerly *Methanosaeta*)*, Unclassified Ignavibacteriales, Spirochaetaceae, Holophagaceae* and *Anaerolineaceae* (Table 2). This suggests that not only the Hg^(II)^ methylators themselves, but also the supporting and interacting bacterial communities residing in the soil environment may influence MeHg formation across the studied regions. *Anaerolineaceae, Spirochaetaceae* and *Holophagaceae* are for example known to generate acetate by fermentation processes (58). *Fibrobacterales*, have recently been suggested to have an important role in cellulose hydrolysis in anaerobic environments, including soils (59). The *Ignavibacteria* class was recently described (Iino et al., 2010) and the physiology and metabolic capacities of this group is still poorly known, even if a distinctive feature of this group is the ability to grow on cellulose and its derivatives with the utilization of Fe(III) oxide as electron acceptor (60). It may well be that these families, which correlated well with %MeHg (Table 2) and seem to be involved in the degradation of long chain OM compounds (61, 62), promoted MeHg production by providing appropriate substrates (e.g. acetate) for the Hg^(II)^ methylators. Hg^(II)^ methylators and non-Hg^(II)^ methylating members of *Desulfobulbaceae*, known to oxidise organic substrates incompletely to acetate (63), might also have provided the necessary substrate to Hg^(II)^ methylators (Table 2). Based on our results, we propose an important role of also the non-Hg^(II)^ methylating bacterial heterotrophs in sustaining the activity of the Hg^(II)^ methylating microorganisms and thereby influencing MeHg formation in boreal forest soils. Moreover, the correlation between *Methanothrix* and %MeHg deserves special attention. It has been shown that *Methanothrix* can establish syntrophic cooperation with *Anaerolineaceae* (61) or *Geobacteraceae* (64) in methanogenic degradation of long chain carbon compounds (alkanes). As our results show that *Geobacteraceae* are major contributors to the Hg^(II)^ methylating microbial community (Table 3), the high correlation found between *Methanothrix* and %MeHg could be the result of the interaction between the non-Hg^(II)^ methylating *Methanothrix* and the Hg^(II)^ methylating *Geobacteraceae.* In brief, we provide novel system-level information on putative trophic interactions between non-Hg^(II)^ methylating and the Hg^(II)^ methylating taxa. We further suggest that more in depth studies with metagenome-level sequencing and metabolic pathway reconstruction will be a logical next step to gain a more complete understanding of how Hg^(II)^ methylating bacterial and archaeal species interact in soils.

## CONCLUSIONS

A newly developed strategy that combine high-throughput *hgcA* amplicon sequencing with molecular barcoding revealed diverse clades of Hg^(II)^ methylators in forest soils. This study confirms a predominant role of *Deltaproteobacteria*, and in particular *Geobacteraceae,* as key Hg^(II)^ methylators in boreal forest soils. *Firmicutes*, and in particular *Ruminococcaceae*, were also abundant members of the Hg^(II)^ methylating microbial community. Besides the identified Hg^(II)^ methylators, we suggest that the non-Hg^(II)^-methylating bacterial community (e.g. *Anaerolineaceae, Holophagaceae* and *Spirochaetaceae*) might have contributed to the net MeHg formation (%MeHg) by processing OM and thereby providing low OM compounds as a substrate to Hg^(II)^ methylators (e.g acetate). By revealing linkages between Hg^(II)^ methylators and non-Hg^(II)^ methylators, our results calls for further community-level work on the metabolic interactions in soil microbial communities to understand Hg^(II)^ methylation. Such studies would need to go beyond the Hg^(II)^ methylating microbial populations. Our findings provide a better understanding of Hg^(II)^ methylating microbial communities in forest soils and the boreal landscape.

## MATERIALS AND METHODS

### Site description

Soil samples were collected from 200 sites in October 2012 and were distributed across eight catchments in three boreal forest regions in Sweden (Table S1 and S2). Within each of the catchments, 25 samples were collected. The most southern region Örebro (59°10´16.39˝N 14°34´3.01˝E) includes three catchments and the sampled soils are dominantly Podzol with Histosols (65) in the lower parts of the catchments along the streams. The organic matter (O) horizons were most often thicker than 20 cm. More detailed information is given in Eklöf et al. (66). Two northern regions, Balsjö (64°1´37˝N 18°55´43˝E) and Strömsjöliden (64°6´48˝N 19°7´36˝E), are located 600–700 km north of Örebro and around 14 km apart from each other. Balsjö includes three catchments dominated by orthic Podzol, with Histosols along the streams. The O horizons were most often thicker than 10–20 cm in the lower parts and less than 10 cm higher up in the catchments. More details are given in Löfgren *et al*. (2009). Strömsjöliden includes two catchments and the soils are dominated by fine-grained moraine. The organic layers are most often less than a few centimetres deep. The samples with high soil MeHg concentrations and %MeHg > 1% were defined as “MeHg hotspots” (n=34), see a summary of the soil characteristics of “MeHg hotspots” in Table S4.

The daily mean air temperatures during the 9 sampling days in September in 2012 varied between 7 and 12 °C in Örebro catchments and 4 and 11 °C in Balsjö and Strömsjöliden catchments. There were no major rain events during the sampling period and the temperature and precipitation was normal for the time of the year.

### Soil sampling

Soil samples were collected with a soil coring tube (Ø=23 mm). In each catchment, around half of the samples (n=12) were collected systematically along the topographic fall line of the hill slope, at set distances from the stream draining the area. These samples were collected from the upper 6 cm of the O horizons or the whole O horizons if these were less than 6 cm deep. The locations of the remaining sampling sites (n=13) were chosen by actively looking for potential hot spots for MeHg formation, such as wet patches, driving tracks and stump holes. These targeted samples were also collected from various depths, e.g. depths where groundwater levels were most frequently fluctuating were of special interest for potential Hg^(II)^ methylation.

Single-use plastic gloves were used and soil samples for chemical analyses were collected in plastic bags or acid washed Falcon tubes and stored on ice in a cooler during transport to the laboratory (within 8 hours). Soil samples for molecular analyses were collected following adequate aseptic sampling protocols. All sampling equipment was sterilized by washing in 70% ethanol in between samples. Samples were collected in sterilized plastic tubes and frozen in liquid nitrogen directly in the field, and then stored at −80°C until further processing and analyses.

### Chemical analyses

Soil samples were analysed for total Hg (THg), MeHg, water content, and mass percentage of carbon (C), nitrogen (N) and sulphur (S). Samples were freeze-dried and ground by hand in a mortar prior to analyses for THg, C%, N% and S%. Wet and dry weights were measured to estimate the water content. Total Hg was measured using a Perkin Elmer SMS100 total Hg analyser in accordance with US EPA method 7473. The method includes a thermal decomposition step, followed by amalgamation and atomic absorption spectrophotometric detection (working range 0.05–600 ng). Reproducibility and accuracy of measurements were checked by analyses of replicate samples and reference standards. Analyses of MeHg were done by using GC-ICPMS (68) on fresh samples immediately after thawing. C, N and S were analysed on dry soils packed tightly in tin capsules (Elemental Microanalysis, 6.4 mm) and subsequently measured by high temperature catalytic oxidation with a COTECH ECS 4010 elemental analyser calibrated with sulfanilamide standard (C 41.84 %, N 16.27 %, H 4.68 %, O 18.58 %, S 18.62 %). Analytical precision was < ± 0.3 % for C, ± 1.5 % for N and ± 3.5 % for S.

### Microbiological analyses

#### 16S rRNA gene

Microbial DNA was extracted from soil samples using the Power soil DNA isolation Kit (MoBio Laboratories Inc, CA, USA) and the quality of the extracted DNA was assessed by gel electrophoresis (1% agarose). Bacterial 16S rRNA genes were amplified in two steps polymerase chain reaction (PCR) according to the protocol in Sinclair *et al* (2015). Briefly, non-barcoded primers Bakt_341F and Bakt_805R (Table S7) were used for the 1^st^ PCR step of 20 cycles. The resulting PCR products were diluted 100 times before being used as template in a 2^nd^ PCR step of 10 cycles with similar primers carrying sample-specific 7-base DNA barcodes. All PCRs were conducted in 20 µL volume using 1.0 U Q5 high fidelity DNA polymerase (NEB, UK), 0.25 µM primers, 0.2 mM dNTP mix, and 0.4 µg bovine serum albumin. The thermal program consisted of an initial 95 °C denaturation step for 5 min, a cycling program of 95 °C for 40 seconds, 53 °C for 40 seconds, 72 °C for 60 seconds and a final elongation step at 72 °C for 7 minutes. Amplicons from the 2^nd^ PCR were purified using the Qiagen gel purification kit (Qiagen, Germany) and quantified using a fluorescence-based DNA quantitation kit (PicoGreen, Invitrogen). The final amplicons after two PCR steps were pooled in equal proportions to obtain a similar number of sequencing reads per sample. Amplicon sequencing was carried out following the protocol described in Sinclair *et al* (2015) using the MiSeq instrument. Illumina sequencing was performed by the SNP/SEQ SciLifeLab facility hosted by Uppsala University using 300bp chemistry. Chimera identification and OTU (Operational Taxonomic Unit) clustering by denoising was done using UNOISE (from USEARCH version 9, ref. 70, 71). SINTAX (from USEARCH version 9, ref. (72)) with the SILVA reference database (release 128) was used as a base to taxonomically annotate OTUs. The sequence data has been deposited to the EBI Archive under accession number PRJEB20882.

#### *HgcA* gene

Among the 50 samples selected based on having %MeHg >1 %, 34 resulted in positive PCR amplification of the *hgcA* gene. The protein-coding gene *hgcA* which plays an essential role in Hg methylation was amplified with previously published *hgcA* primers (*hgcA_ 261F* and *hgcA_912R*) (Table S7, 34) modified for parallelized high-throughput Illumina sequencing. HPLC-purified primers carrying Illumina adaptors at the 5’ end (*hgcA_261F_Adaptor* and *hgcA_912R_Adaptor*, Table S7) were here used for the 1^st^ stage PCR. In the 2^nd^ stage PCR, standard Illumina handles and barcode primers (Table S7) were used to enable pooling of all the samples for parallelized Illumina sequencing. *HgcA* was first amplified in 50 µL volume with 1x Phusion GC Buffer, 0.2 mM dNTP mix, 5% DMSO, 0.1 µM of each adaptor-linked primer, 7 µg/µL BSA, 4 µL extracted DNA template, and 1.0 U Phusion high fidelity DNA polymerase (NEB, UK) for an initial denaturation of 2 min at 98 °C followed by 35 cycles (10 s at 96 °C, 30 s 56.5 °C and 45 s at 72 °C), and a final extension at 72°C for 7 min. Following this initial step, a 2^nd^ PCR was conducted to add sample-specific molecular barcodes. Reactions were carried out in 20 µL volumes using 1x Q5 reaction buffer, 0.2 mM dNTP mix, 0.1 µM barcode primers, purified 1^st^ PCR products and 1.0 U Q5 high fidelity DNA polymerase (NEB, UK) for an initial denaturation of 30 s at 98 °C followed by 18 cycles (10 s at 98 °C, 30 s 66 °C and 30 s at 72 °C), and a final extension at 72°C for 2 min. The quality and size of the *hgcA* amplicons were assessed by gel electrophoresis and GelRed visualization on a 1% agarose gel (Invitrogen, USA) prior to purification by Agencourt AMPure XP (Beckman Coulter, USA) after both PCR steps. Quantifications of purified amplicons from the 2^nd^ stag PCR were performed using the PicoGreen kit (Invitrogen).

Amplicons were sequenced using the same method as for the 16S rRNA gene. Forward read sequences were only used in data analysis due to long PCR product. Low quality sequences were filtered and trimmed using SICKLE (73) and adapter were removed by using CUTADAPT (74). Subsequent processing of reads were performed by USEARCH and clustered at 60% identity cutoff using cd-hit-est (75). HMMER (76) search was used for taxonomical annotation with manually curated database of *Proteobacteria* and sequences of Podar *et al.* (2015). More details can be found in Bravo *et al.* (2018).

#### Phylogenetic analysis

A phylogenetic analysis was performed for *hgcA* sequences representative for the OTUs observed for the 34 hotspots and existing *hgcA* entries in our curated database. The sequences were adequately curated and taxonomy homogenized using taxtastic [https://github.com/fhcrc/taxtastic] and the R-package taxize (77). The obtained protein sequences were aligned with MUSCLE (78) (version 3.8.1551). The alignment was trimmed to the size of the amplicon, and a tree was generated using RAxML (79) (version 8.2.4) - with the PROTGAMMLG model and autoMR to choose the number of necessary bootstrap resamplings (n = 750). This tree and the corresponding alignment were used to generate a reference package for PPLACER (80). The guppy tool of PPLACER was then used to classify the sequences with a likelihood threshold of 0.8.

### Statistical analysis

Family-level microbial community composition in the different samples were compared using non-metric multidimensional scaling (nMDS) based on Bray-Curtis similarities and using the software PRIMER 7 (81). Information on the common set of samples from community composition based on Bray-Curtis similarities and that from geochemical variables based on Euclidean distance was presented in one single ordination. A combined nMDS plot with bubble and vector plots of geochemical factors projected on the same ordination of community composition was constructed to reveal the relationships between community compositions and potentially explanatory geochemical variables (81). Pearson’s correlation coefficient (R) was assessed to reveal linear relationships between variables using a significance level of alpha < 0.05.

## Acknowledgements

This project was carried out within the Swedish-Sino SMaREF (2013-6978) funded by the Swedish Research Council. This study was also supported by the Swedish Energy Agency (grant number 36155-1) and the Swedish Research Council (Grants 2011–7192 and 2012-3892) and Generalitat de Catalunya (Beatriu de Pinos BP-00385-2016). Sequencing was carried out at the SciLifeLab SNP/SEQ facility hosted by Uppsala University and we also acknowledge the Uppsala Multidisciplinary Centre for Advanced Computational Science (UPPMAX) for access to storage and computational resources.

